# Mechanisms of Anammox Adaptation to High Temperatures: Increased Cyclization of Ladderane Lipids and Proteomic Insights

**DOI:** 10.1101/2024.07.23.604647

**Authors:** Karmann Christina, Navrátilová Klára, Behner Adam, Noor Tayyaba, Danner Stella, Majchrzak Anastasia, Šantrůček Jiří, Podzimek Tomáš, Marin Lopez Marco A., Hajšlová Jana, Lipovová Petra, Bartáček Jan, Kouba Vojtěch

## Abstract

Graphical abstract

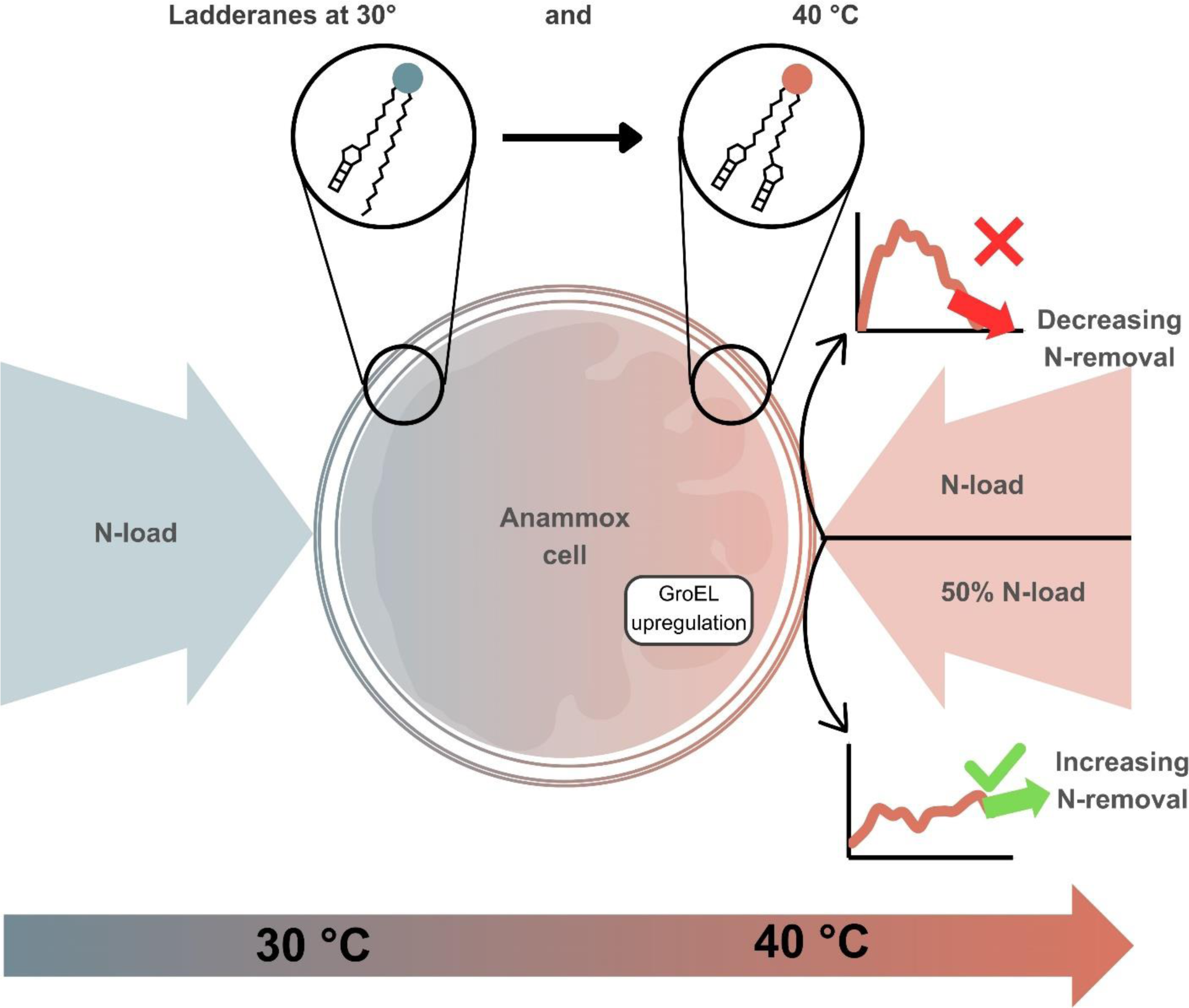

Although anammox-based processes have been widely applied in mesophilic conditions of reject water and recently in mainstream conditions, the potential of their implementation in high-temperature wastewaters remains largely unexplored. Therefore, this study investigated the operation parameters for the successful adaptation of anammox bacteria and the mechanisms involved on the proteomic and cellular level including unique ladderane lipids. For this purpose, the enrichment of ‘*Candidatus* Brocadia’ was cultivated in two fed-batch reactors (FBRs) at a lab scale. The temperature of one FBR was gradually increased from 30 to 40 °C while the other FBR was maintained at 30 °C with four consecutive replicates of this experiment. For this adaptation to be successful, the original loading rate had to be at least halved, or ideally maintained below half the value of the specific anammox activity at the time. The most notable adaptation mechanisms included: (1) upregulation of chaperones and (2) doubled ladderane cyclization via the replacement of non-ladderane fatty acid by a ladderane fatty acid in ladderane lipids (p-value 0.005). To our best knowledge, this is the first study to describe the novel mechanism of ladderane cyclization which together with other adaptation strategies presents crucial indicators in anammox adaptation to high-temperature wastewaters.

## 1 Introduction

Anaerobic ammonium-oxidation (anammox) is a microbial process used for nitrogen removal in wastewater treatment plants. Nowadays, anammox bacteria are commonly used to treat the anaerobic digestion effluent and industrial wastewaters at mesophilic temperatures (30-35 °C). The lower sludge production, limited need for oxygen, and complete lack of carbon source requirements led to the recent adoption of anammox process at full scale in mainstream conditions (10-25 °C) of municipal wastewater treatment plants (Fofana et al., 2022; Yuan et al., 2021). However, a large amount of nitrogenous wastewater is produced from the thermophilic anaerobic digestion of sludge or various high-temperature industrial wastewaters such as thermomechanical pulping wastewater or streams from hydrothermal carbonisation surpassing the temperature of 35 °C.

Recent studies addressed the issue of thermophilic anammox coined in a range from 35 to 55 °C by Sobotka et al. (2016) at lab-scale conditions (Vandekerckhove et al., 2020; Zhang et al., 2018). They reported rapid system deterioration above 40 °C, accompanied by a decreased nitrogen removal rate (NRR) and loss of activity. Vandekerckhove et al. (2020) were able to maintain reactor operation even at 40 °C, resulting in the slow adaptation of anammox to 50 °C and an increase in activity. Moreover, in nature, anammox bacteria were found to be able to survive at temperatures as high as 80 °C (Byrne et al., 2009). However, so far, it is unclear what mechanisms the anammox bacteria use for adaptation to high temperatures.

Microorganisms, including anammox, can adapt to different temperatures by several mechanisms, including changes at proteomic and cellular levels, in transcriptome, and genome, but also on the level of changes in microbial population. The crucial aspect of the response to thermophilic conditions at the proteomic level is stabilizing the natural protein configuration via thermoprotectant enzymes and facilitating correct protein folding (Wang et al., 2023). The response at a cellular level commonly involves a homeoviscous adaptation of membranes. This process maintains optimal membrane fluidity by altering lipid structure to prevent energy loss. At increasing temperatures, bacteria produce a combination of more branched fatty acids (FAs), FAs with longer chains, and FAs with more double bonds and less disruptive smaller head groups (Jebbar et al., 2015). Furthermore, anammox bacteria are generally considered to be the only organisms containing the so-called ladderane lipids apart from standard lipids. Ladderanes consist of a glycerol backbone with *sn-3* position occupied by a polar head group (phosphotidylcholine (PC), phosphatidylethanolamine (PE) or phosphatidylglycerol (PG)). Their *sn-2* position is occupied by alcohol with three concatenated cyclobutane rings and one cyclohexane [3]-ladderane, specifically C20-[3]-ladderane. The *sn-1* position may contain any normal alkyl or a ladderane FA with 18 to 20 carbons and five cyclobutane rings ([5]-ladderanes) or [3]-ladderane, although ladderane fatty acids with 16, 22 and 24 carbons have also been identified (Rattray et al., 2010). Some mechanisms of ladderane alteration (ladderane and lipid shortening, different polar head group) and also changes at proteome level (chaperones, cryoprotectants and cross-feeding pathways) were well described for anammox adaptation to lower temperatures (Kouba et al., 2022a; Kouba et al., 2022c; Rattray et al., 2010).

Although anammox-based processes may present a solution for sustainable nitrogen removal from high-temperature wastewaters, the mechanisms of anammox adaptation to thermophilic range and suitable regime for achieving such adaptation remain undescribed. Therefore, this study investigated the suitable operation conditions for achieving anammox adaptation and also mechanisms involved in this process including changes in ladderanes structure and proteome.

## 2 Materials and Methods

### 2.1 Inoculum

Floccular biomass enriched in ‘*Candidatus* Brocadia’ was adopted as inoculum from a cultivation reactor from Radboud University in Nijmegen (Netherlands). This fed-batch reactor (FBR) was operated for more than 100 days, at 30 °C and a pH of 7.2, which was automatically adjusted using the application LabView (National Instruments, Texas, USA) by addition of 1M KHCO_3_ (pH buffer). The anoxic conditions were maintained by sparging the liquid with a mixture of N_2_/CO_2_ (95%/ 5%), the composition of the synthetic substrate used for feeding the bioreactors is shown in Table 1 in the Supplementary materials, the nitrogen was added in the form of NaNO_2_ and NH_4_Cl in different concentrations depending on the concentration of biomass and other parameters. The cultivation reactor was sampled for inoculum of all four experiments and immediately transferred into experimental reactors.

### 2.2 Reactor operation

Two FBRs (R0 and R1) with working volumes of 1 L were operated as the cultivation reactors in three consecutive experiments. Their scheme is shown in Figure 1 and a photograph in Supplementary materials, Fig. 1. Each cycle lasted 8 hours consisting of reaction time (445 min), settling (25 min) and effluent draw (10 min). This cycle was performed with timers and real-time controller system (RSC) CompactRIO® and the application LabView (National Instruments, Texas, USA). FBRs were fed with the same synthetic substrate as the cultivation reactor.

**Figure 1:**
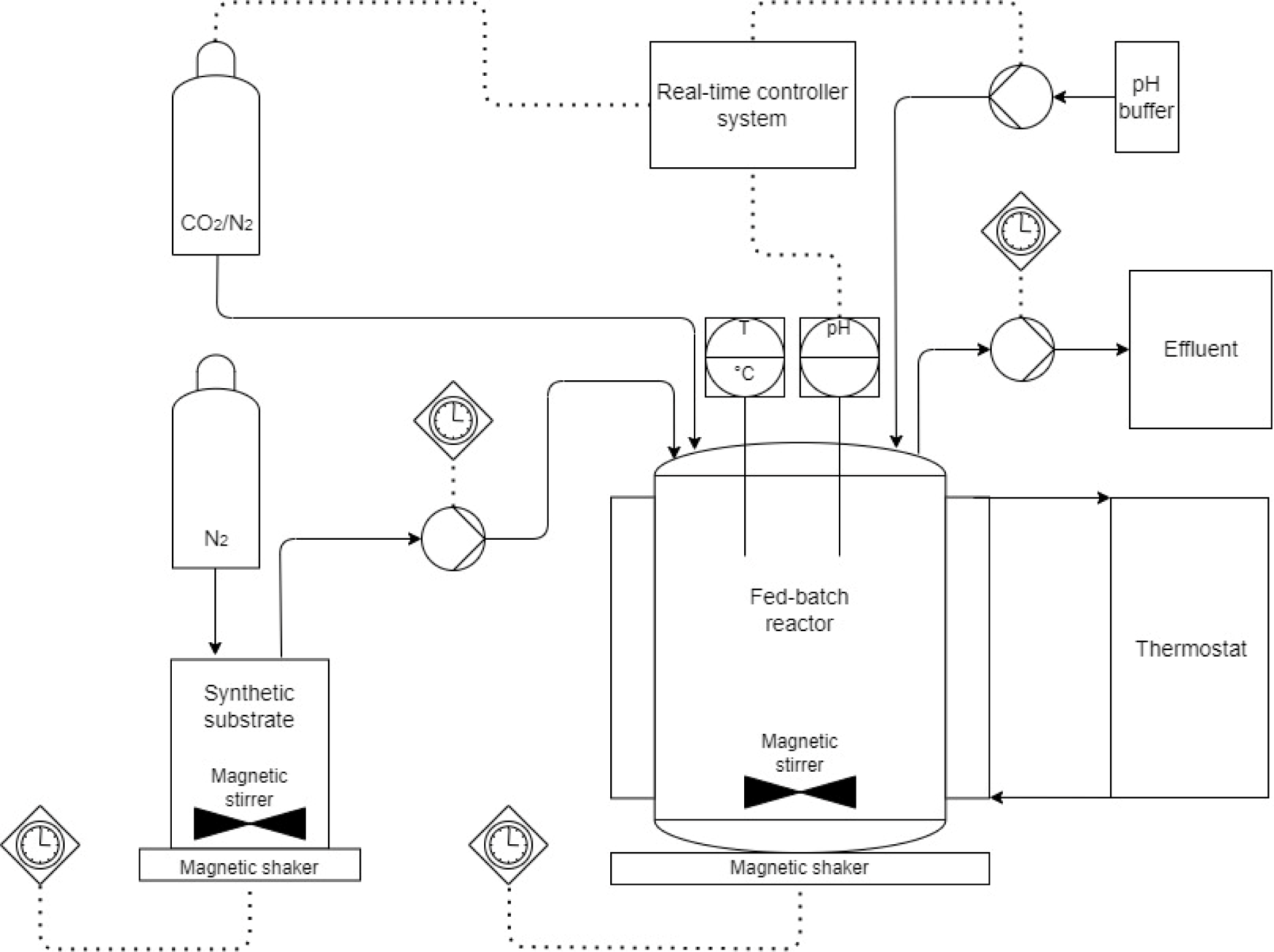
Scheme of the reactors used for the adaptation of anammox enrichments to a high temperature.

The temperature of the control reactors was maintained at 30 °C as the cultivation reactor. Immediately after inoculation, the R1 was subjected to a temperature increase to more than 35 °C and then progressively to 40 °C over the course of 40 to 60 days and maintained for a minimum of 5 days before further analysis and biomass sampling.

Due to the molar ratio of NH_4_^+^ and NO_2_^-^ of 1:1, there was always a surplus of NH_4_^+^, preventing NO_2_^-^ poisoning. A complete NO_2_^-^ removal was aimed for and in case of NO_2_^-^ accumulation, nitrogen loading rate was reduced (NLR).

The substrate and effluent of reactors were regularly sampled weekly to determine concentrations of NH_4+_-N, NO_3-_-N, and NO_2-_-N measured using Gallery™ Discrete Analyzer (Thermo Fisher Scientific, Waltham, USA) according to the standard methods (Apha, 1998). NLR (kg-N/m^3^/d), NRR (kg-N/m^3^/d), and Nitrogen removal efficiency (NRE, %) were calculated by Eqs. (1-3) (He et al., 2018). Furthermore, colourimetric measurement via Nitrite Test MQuant™ 2-80 mg/l NO_2_ ^-^ test stripes (Merck) was used for the day-to-day analysis of the performance of the bioreactors and ability to sufficiently degrade influent nitrogen avoiding harmful nitrite poisoning. Biomass concentration was measured as total suspended solids (TSS) and volatile suspended solids (VSS) according to standard methods (Apha, 1998). NLR and NRR were related to VSS, resulting in the Bimass-specific nitrogen loading rate (SLR) and Biomass-specific nitrogen removal rate (SRR), respectively, with units kg-N/kg-VSS/d.

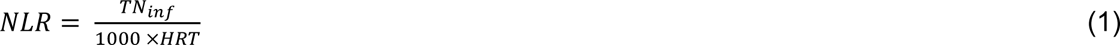

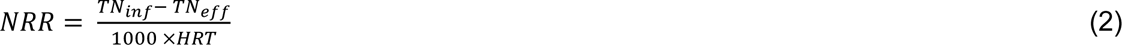

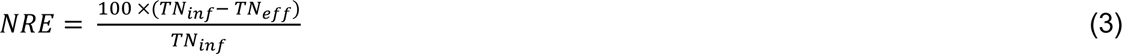

### 2.3 Specific anammox activity

Anammox activity was determined by *in-situ* specific activity tests at the beginning of the experiment after inoculation (30 °), after reaching 39 °C and at the end of the experiment (40 °C) before the biomass was harvested for further analysis. The reactors were spiked with NH_4_ ^+^-N and NO_2_ ^-^-N in a 1,1:1 molar ratio. The reactors were then sampled every 15 min, the samples were filtered through standard 0.45 µm glass fibre filters and analyzed spectrophotometrically on the Gallery™ Discrete Analyzer (Thermo Fisher Scientific) according to the standard methods (Apha, 1998). SAA was measured as the sum of NH_4_ ^+^-N and NO_2_ ^-^-N removal rates per biomass concentration and time. The nitrogen concentration change over time was fitted with linear regressions, whose slopes were determined as volumetric removal rates according to Kouba et al. (2022c). The activities were subsequently used for the determination of activation energies (Ea) following the Arrhenius empirical model shown in Eqs. (4-5). The ‘k’ represents the ratio of anammox activities at lower (numerator) and higher (denominator) temperatures, ‘ln’ denotes the natural logarithm, ‘A’ is a constant pre-exponential factor, ‘Ea’ signifies the activation energy (J/mol), ‘R’ stands for the ideal gas constant (J/mol/K), and ‘T’ represents the thermodynamic temperature (K). Afterwards, the dataset underwent linearization using equation (5).

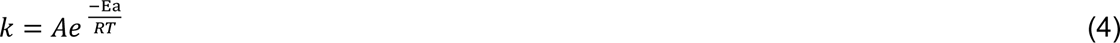

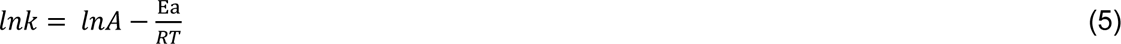

### 2.4 Microbial analysis

The biomasses at the beginning and at the end of experiments were sampled from R0 and R1 for the analysis of microbial community and lyophilized. Firstly, 250 μg of lyophilized biomass was harvested for DNA extraction using DNeasy® Power-Soil® Kit (Qiagen GmBH, Germany), its purity was verified via BioDrop μLite machine (BioDrop Ltd, UK) and concentration was analyzed by Qubit® Fluorometer with Qubit^TM^ dsDNA HS Assay Kit.

Subsequently, a relative qPCR assay using anammox-specific 16S rRNA and general 16S rRNA genes was performed with KAPA SYBR® FAST qPCR Master Mix (2X) Kit (Kapa Biosystems, Inc.). The primers used for the amplification of anammox-specific 16S rRNA genes (Anammox 16S) were: 5′-GTCRGGAGTTADGAAATG-3′ (A438f) and the reverse 5′ - ACCAGAAGTTCCACTCTC-3′ (A684r) while the universal primers (Universal 16S) used were the 5′-CGGCAACGAGCGCAACCC-3′ (1114f) and 5′-CCATTGTAGCACGTGTGTAGCC-3′ (1275r). qPCR analysis was performed using a C1000™ Thermal Cycler (BioRad Laboratories Inc., USA). Initial activation for both assays was maintained at 95 °C for 5 min. The reaction conditions for the Anammox 16S were as follows: 35 cycles of 5 s at 95 °C (denaturation), 15 s at 55 °C (annealing), and 15 s at 72 °C (elongation). Similarly, qPCR parameters for Universal 16S followed these steps: 30 cycles of 5 s at 95 °C, 10 s at 51 °C, and 15 s at 72 °C. The relative quantification was determined mathematically in terms of the relative expression ratio in qPCR, developed by Pfaffl (2001).

Furthermore, the results were confirmed via amplicon sequencing. A two-step PCR targeting the V4 and V5 regions of the 16S rRNA gene was used for libraries preparation via the Nextera technology. The primers used for the amplicon sequencing were 515F (5’-GTGYCAGCMGCCGCGGTAA-3’) and 926R (5’-CCGYCAATTYMTTTRAGTTT-3’). A mock community, which was composed of 15 bacterial strains, was included as a positive control (Fraraccio et al., 2017). Libraries were sequenced using the NovaSeq 6000 system.

### 2.5 Proteomics

#### 2.5.1 LC-MS/analysis

The biomass of each sample was disrupted using OneShot System (Constant Systems, UK). Amount of proteins was set to 100 After that, the samples were prepared using EasyPep™ MS Sample Prep Kit (ThermoFisher Scientific) according to the protocol. Mass spectra were acquired using a Maxis Impact ESI-QToF mass spectrometer (Bruker Daltonics, Bremen, Germany) connected with a Dionex Ultimate3000 RSLCnano UHPLC chromatograph (Thermo Scientific™, Waltham, MA, USA). Dry samples were resuspended in 50 µl of 3% v/v acetonitrile and 0.1% v/v formic acid (peptide concentration approximately 2 µg/µl). Subsequently, 1 µl of the sample was loaded on an Acclaim PepMap 100 trap column (100 µm × 2 cm, C18 reversed phase, particle size: 5 µm; Thermo Scientific) using a flow rate of 7 µL/min. After 5 min of washing, the flow was directed to an Acclaim PepMap RSLC C18 analytical column (75 µm × 150 mm, C18 reversed phase, particle size: 2 µm; Thermo Scientific). The mobile phase A was 0.1% v/v formic acid in water and the mobile phase B was 0.1% v/v formic acid in acetonitrile. Peptides were eluted with a linear gradient of 3–35% B for 180 min followed by column washing (90% B) and re-equilibration of the columns (3% B) before loading the next sample. The flow rate during the peptide elution was set to 0.3 µL/min. The eluted peptides were introduced directly into a Captive spray ESI source (Bruker Daltonics, Bremen, Germany). The spray voltage was set to 1 200 V, the temperature to 150 °C, and the flow of dry nitrogen to 3 L/min. Measurements were performed in the data-dependent analysis mode. Mass spectra were recorded every 3 s in the range of 50–2200 m/z. Precursors for collision-induced dissociation were selected in the range of 400–1400 m/z and MS/MS spectra were collected at the speed of 4–16 Hz depending on the intensity of the precursors. Nitrogen was used as a collision gas. The preferred charge range was 2–5, with single-charged precursors being excluded; a dynamic exclusion time of one minute was applied. Both profile and line spectra were saved.

#### 2.5.2 Data processing

Raw data were processed with MaxQuant version 1.6.10.43 using the Andromeda search engine for protein identification. The following parameters were set: carbamidomethylation of cysteines as a fixed modification, protein N-terminal acetylation and methionine oxidation as variable modifications, and enzyme trypsin with two missed cleavages allowed. The spectra were searched against the ‘*Candidatus* Brocadia fulgida’ database downloaded from the Uniprot web page on July 01, 2022, combined with a database of common laboratory contaminants, the reversed database was used for false discovery calculation. For precursors, a tolerance of 0.07 Da for the first search and 0.006 Da for the main search was set. Fragment ions were matched with a tolerance of 40 ppm. Match between runs was turned on with a match time window of 2 min set. Peptide and protein identification were filtered so that a 1 % false discovery rate was maintained. Label-free quantification was performed using MaxLFQ algorithm. LFQ ration min. count was set to 1, other parameters were left on their default values.

Statistical analysis was performed using Perseus version 2.0.9.0 using proteingroups.txt file from MaxQuant as input. First, protein groups labelled as Potential contaminants, Reverse and Only identified by site were removed. Samples were assigned into one of the three groups based on the cultivation temperature – inoculum, 30 and 40 °C. The LFQ values were log2 transformed and then protein groups were filtered based on valid LFQ values (at least two valid values in each of the three groups). Next, one-way ANOVA was used to assess the statistical significance of protein abundances using a 10 % permutation-based FDR adjustment. Finally, post hoc Tukey’s HSD was performed with an FDR of 10%.

### 2.6 Ladderane analysis

The samples for ladderane analysis were sampled at the end of each Run after at least 5 days of operation of R1 at 40 °C and simultaneously from R0 (30 °C) with a dried weight of 0.3 g. The sample preparation and analysis were done according to Kouba et al. (2022b) by lyophilizing the samples and subsequently shaking them in an extraction solvent mixture (MeOH: DCM, 2:1, v/v). Afterwards, the samples were sonicated, centrifuged (5 min, 10,000 rpm, 5 °C) and filtered through a 90 μm filter paper, grade 390 (Ahlstrom, Munksjö). The resulting supernatants were analyzed by the Dionex UltiMate 3000 RS ultra-high performance liquid chromatography (U-HPLC) system (Thermo Fischer Scientific, Waltham, USA), coupled to quadrupole-time-of-flight SCIEX TripleTOF® 6600 mass spectrometer (SCIEX, Concord, Ontario, Canada). The data was processed based on Hurkova et al. (2019). The specific amount of each ladderane structure was related to their total quantity or the quantity of a subgroup. The statistical significance of differences between samples from R0 and R1 was verified by analysis of variance with a p-value below 0.05.

## 3 Results and discussion

### 3.1 Performance of anammox reactors at increasing temperatures

The inoculum cultivated at 30 °C was used for starting R0 and R1. While the temperature of R0 was maintained at 30 °C, the temperature of R1 was gradually increased from 30 to 40 °C (Fig. 2). On the second day of the experiment, the temperature of R1 was elevated to 35 or even to 38.5 °C and then progressively brought up to 40 °C.

**Figure 2:**
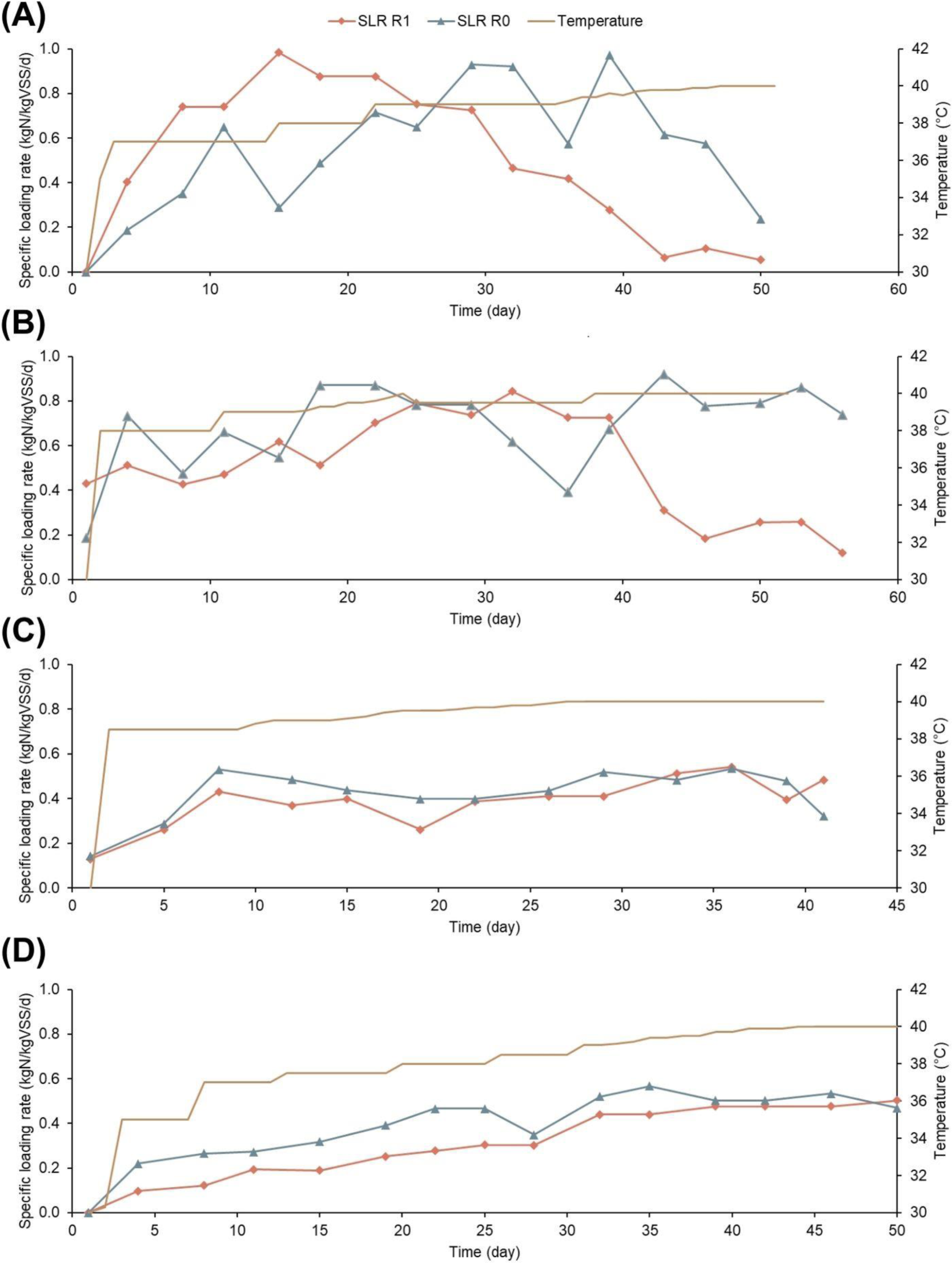
Development of specific loading rate of R0 and R1 and temperature of R1 in the four Runs. A) Run 1, B) Run 2, C) Run 3, D) Run 4.

SLR was used as a determining parameter to account for the biomass concentration, which varied from 0.6 to 4 kg-VSS/m^3^ among Runs 1-4, based on our analysis. Therefore, SLR makes the comparison of different Runs clearer and its development in different Runs is depicted in Fig. 2. SLR of R1 was raised as high as 1.0 kg-N/kg-VSS/d (corresponding to SRR of 0.8 kg-N/kg-VSS/d) in Run 1 and 2 and then decreased below 0.1 kg-N/kg-VSS/d due to repeated NO_2_ ^-^ accumulations followed by a decrease in removal rate and NRE. The SLR of R0 in Run 1 also had to be decreased, however, due to operational issues with pH adjustment. On the other hand, R0 in Run 2 was repeatedly exposed to similarly high loading without a resulting major decline. Correspondingly to the development of SLR, the NREs of R1 (Runs 1-2) were 84 and 85 % in the beginning, comparable to R0 with NREs of 84 and 87%. However, when the temperature exceeded 39°C, the NREs dropped close to 70 %. Therefore, another two runs (Runs 3-4) were performed with a lower SLR.

Zhang et al. (2018) observed a significant decrease of NRR at 40 °C from 1.5 to 0.25 kg-N/L/d even though the inoculum was cultivated at 35 °C (a difference of 5 °C), while the noticeable decrease in this study occurred only after a change of more than 9°C This suggests that the temperature of 39-40 °C is a “breakpoint” in the case of anammox bacteria. Similarly, Vandekerckhove et al. (2020) observed a drop in NLR at 43°C from 0.2 to below 0.1 kg-N/L/d, only a 6°C higher temperature than used for inoculum cultivation (37°C). Narita et al. (2017) showed “*Ca.* Brocadia” long-term cultivated at 37°C to lose less activity (16 %), when exposed to 45 °C than the biomass cultivated at room temperature (32 %). Therefore, it is likely that a certain temperature increase, not a specific temperature, provokes an adverse response from anammox bacteria. A similar phenomenon also occurred at decreasing temperatures, with a 10 °C decrease, the NRR declined to half (Gilbert et al., 2014; Isaka et al., 2008).

The median of SRR in all four Runs in this study was 1.0 kg-N/kg-VSS/d but led to a deterioration of the reactors in Runs 1 and 2. Therefore, the maximum SRR was maintained below 0.5 kg-N/kg-VSS/d in Runs 3 and 4 via a slow increase of SLR and resulted in a successful adaptation with stable NREs above 80 %. The maximum SRR in these two runs was comparable to the value of Vandekerckhove et al. (2020), who achieved successful adaptation even at 50 °C. Furthermore, this study reached a similar NRR of up to 1.4 kg-N/L/d as the one at the beginning of the study by Zhang et al. (2018) at the 40 °C phase. This successful adaptation strategy by lowering the SLR suggests a negative cumulative impact of temperature above 39 °C and high SLR.

### 3.2 Development of anammox activity and activation energy

Specific anammox activity (SAA) in both reactors across the four Runs varied from 0.5 to 1.0 kg-N/kg-VSS/d (Fig. 3). The SAAs were the highest at 39°C and notably decreased when the temperature of R1 was brought up to 40 °C and maintained there. Correspondingly, the Ea obtained at temperature range 30-39 °C (−23.8 ± 22.8 kJ.mol^−1^) was notably lower than at 30-40 °C (50.6 ± 36.0 kJ.mol^−1^), shown in Table 2. As shown by Sobotka et al. (2021) who observed a rapid decline of SAA at higher temperatures compared with low-temperature range connected with the impact of heat on proteins and enzymes. Similarly to the findings in this study, Zhu et al. (2017) and Li et al. (2018), who obtained values of 43 and 46 kJ.mol^−1^, respectively, at a 10 °C difference, although at temperatures 25 – 35 °C.

**Figure 3:**
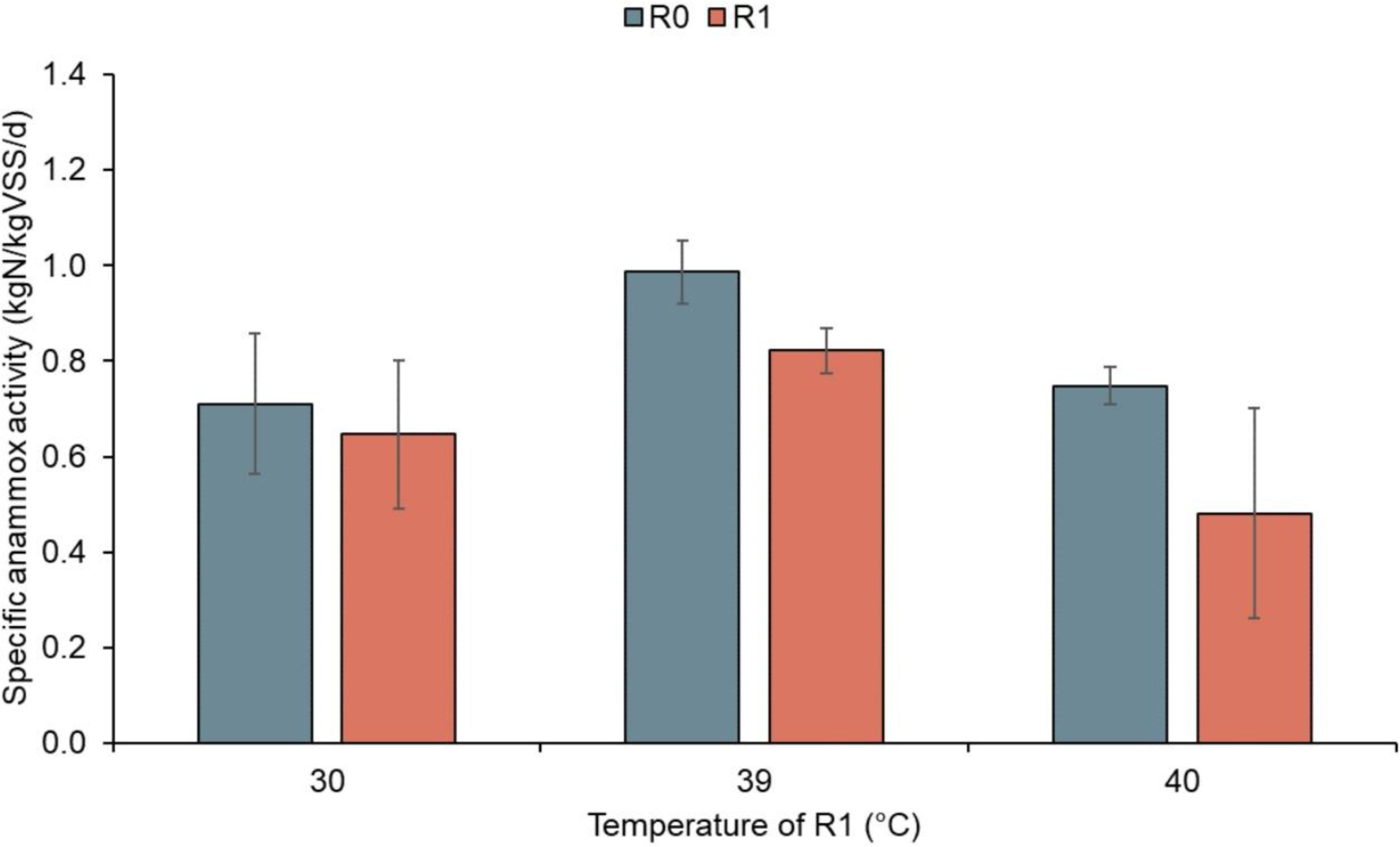
The average SAAs of all four Runs at temperatures 30, 39 and 40 °C.

**Table 2:**
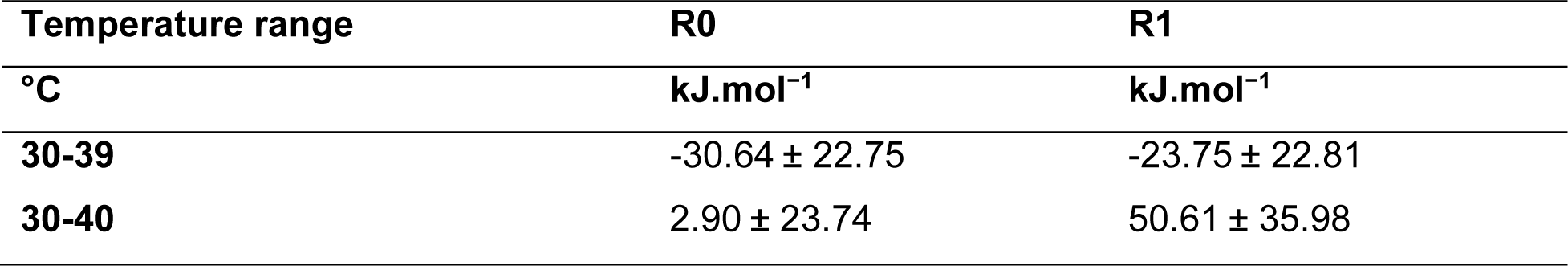
The average activation energies of all four Runs, related to 30 °C.

| Temperature range | R0 | R1 |
| --- | --- | --- |
| °C | $\text{kJ.mol}^{-1}$ | $\text{kJ.mol}^{-1}$ |
| 30-39 | $-30.64 \pm 22.75$ | $-23.75 \pm 22.81$ |
| 30-40 | $2.90 \pm 23.74$ | $50.61 \pm 35.98$ |

The decline occurred particularly in Runs 1 and 2 where the SAA declined to 0.2 and 0.4 kg-N/kg-VSS/d, respectively. Similar decline from 0.5 to 0.2 kg-N/kg-VSS/d was detected by Zhang et al. (2018) when increasing the temperature from 35 to 40 °C. Other studies show SAA reduction to a half or lower when the temperature was brought down or up by 10 °C (Lotti et al., 2015; Park et al., 2017; Sobotka et al., 2016).

These findings are similar to the ones by Zhu et al. (2017), who suggested that the optimum temperature with the maximum SAA is attained at 40 °C. Yet, other studies indicate it may be lower, 37 °C (Narita et al., 2017), 35 °C (Li et al., 2018; Park et al., 2017) or even 30 °C (Lotti et al., 2015; Park et al., 2017) and at higher temperatures, the anammox process may be partially or completely inhibited. However, the temperature at which anammox have the highest activity may be harmful for long-term cultivation.

However, the SAA declined only slightly to 0.8 kg-N/kg-VSS/d in Runs 3 and 4 in both R0 and R1 and this may be therefore attributed to other operation conditions (stirring, DO). The reduction in SLR to half of that of the current SAA led to the retaining of stable SRR and NRE values comparable with the control reactor (R0). Moreover, the SAA was in this way maintained only on a slightly lower level than at 39°C, reaching much higher values than reported by Zhang et al. (2018) and showing the possibility of adaptation.

### 3.3 Changes in microbial community

Samples of inoculum, R0 and R1 were harvested at the beginning or the end of the experiment, respectively, and analysed via NovaSeq sequencing. All the sequences belonging to Planctomycetota phylum were with >99 % similarity identified as a recently discovered ‘*Ca.* Brocadia’ (Narita et al., 2017).

A decline of anammox enrichment (ratio of anammox bacteria to other organisms in the biomass) with temperature was evident. While the anammox population of R0 showed a slight reduction to 8.0 ± 0.3 % compared to inoculum (9.8 ± 0.3 %), it decreased to 3.7 ± 0.3 % in R1. The decrease in R0 abundancy may theoretically be also attributed to factors, such as suboptimal stirring, DO and others. However, the temperature was likely the main factor reducing the anammox abundance at 40 °C to half of the enrichment at 30 °C. Furthermore, this finding is consistent with the decline of nitrogen removal in the R1 reactors, which may be attributed to the decreasing anammox enrichment.

To detect the impact of temperature on anammox activity without regard to the effects of biomass wash-out and abundance reduction, the SAA was related to anammox abundance. The biomass of R1 at 40 °C showed a 130% (± 72 %) activity increase compared to the biomass of R0 at 30 °C. This activity growth was most pronounced for Run 3 in which the R1 activity was 207 % of the R0, while in Run 1 only 65 % of R0. Zhang et al. (2018) detected a loss of NRR, which was found to be directly related to the population of anammox bacteria (Isaka et al., 2008), unlike Vandekerckhove et al. (2020) cultivating anammox in a fixed-bed biofilm reactor (FBBR), which secured their retention and was able to achieve high NLR at 50 °C and the discovery of a novel thermophilic anammox species. In this study, we also observed a reduction in nitrogen removal with increasing temperature. However, its primary cause lay in the decreasing anammox population. Although, a high SLR led to a loss of anammox activity, the SLR reduced to half of the original value increased the anammox activity relative to their abundance. Therefore, adaptation of anammox to high temperatures should mainly focus on anammox wash-out prevention and avoiding an overloading of the system, which results in higher anammox activity and consequently, increased capacity to remove nitrogen in the long-term.

Relative quantification of anammox bacteria to all organisms was performed via qPCR to validate the sequencing results and elucidate a temporal pattern of anammox enrichment development to temperature. The results confirmed the decline of anammox abundance with rising temperature to one-half of the original value. Moreover, the similarity in the anammox enrichment of R1 to R0 (74 %) relative to the inoculum enrichment, was greater when the R1 was operated at 38°C (78 %) than at 40 °C, further implying detrimental impacts of high temperature on the microbial community.

The sequencing also revealed other changes in the microbial population. Obligately anaerobic thermophiles from class Anaerolineae (Nunoura et al., 2013) and other obligately anaerobic bacterium PHOS-HE36 from class Ignavibacteria, also often reported thermophilic (Chen et al., 2023; Iino et al., 2010) increased at 40 °C while the Rhodospirillales and Burkholderiales, genus *Denitratisoma*, associated with facultatively anaerobic or even aerobic conditions decreased (Huang et al., 2019; Pfennig and Trüper, 2019; Ribeiro et al., 2022) further supporting that an excess of DO was not the major contributing factor to abatement of anammox bacteria.

### 3.4 Ladderanes composition

#### 3.4.1 High ladderanes content and increlative reased cyclization as responses to high temperature

The biomass for ladderane analysis was harvested at the end of each Run from both R0 (30 °C) and R1 (40 °C). The quantity of ladderanes (g-ladderanes/ g-lyophilized biomass) was related to anammox relative abundance analyzed via qPCR to its value of inoculum. This relative ladderane content related to anammox abundance (RLC) significantly increased (p-value < 0.05; p-value 0.047). The RLC doubled from 0.8 ± 0.3 in R0 at 30 °C to 1.6 ± 0.1 in R1 at 40 °C.

Furthermore, a significant increase of ladderanes with two ladderane FAs (p-value 0.005) at the expense of ladderanes with a single FA was detected. The structures of these ladderane lipids and fatty acids discussed in this study are presented in Fig. 4.

**Figure 4:**
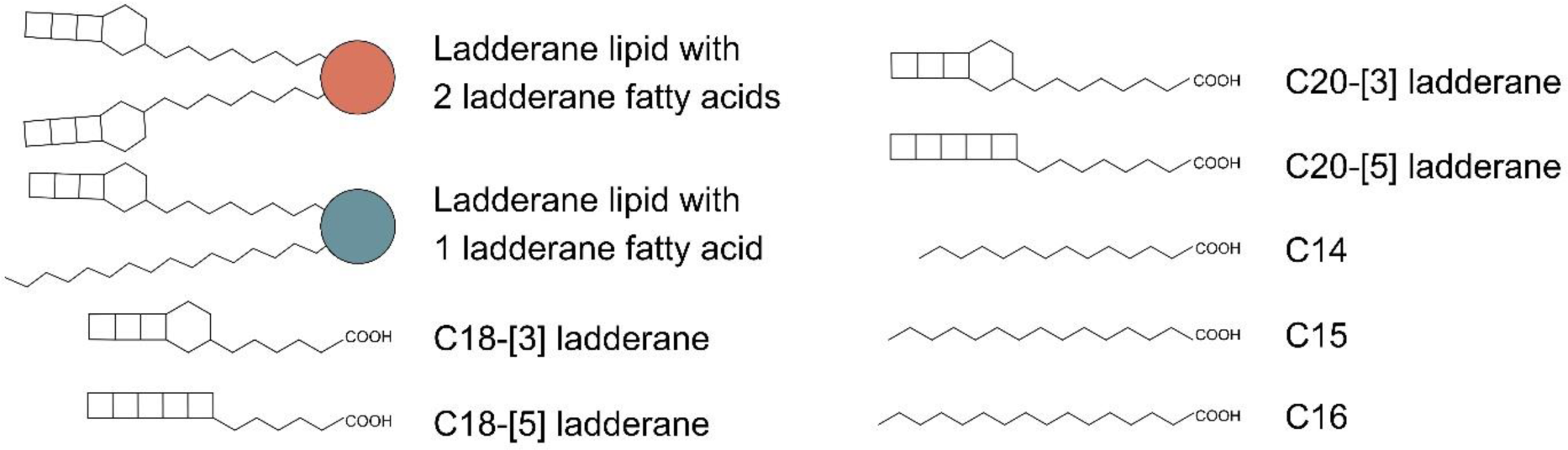
The structures of ladderane lipids and their ladderane and non-ladderane fatty acids (FAs). The non-ladderane FAs (C14, C15 and C16) can also include their isomers. The red or blue circle in the structures of ladderane lipids with one or two ladderane FAs stand for an ester or ether bond to the glycerol backbone and the *sn-3* position occupied by phosphatidylethanolamine, phosphatidylglycerol or phosphatidylcholine.

The relative amount of lipids with ladderane moiety in both positions grew from 0.08±0.01 to 0.14±0.01 (Fig. 5). To the best of our knowledge, we are the first to observe this mechanism, cyclization, and propose it to be an adaptation response to high temperatures. Although ladderanes or cyclobutane rings are not contained in the membranes of any other organism, various bacteria and archaea introduce other cyclic structures into their lipids. For example, cyclopropane-containing FAs made membranes more fluid at low temperatures, however, their rigidity was still higher compared to double bonds (Maiti et al., 2023). However, at sub-freezing temperatures, their ability to disturb the packing order was no longer sufficient and the bacteria used unsaturation instead (Siliakus et al., 2017). Others found the introduction of cyclopropanes to promote membrane fluidity while restricting the transmembrane diffusion of protons and stabilizing membranes against adverse conditions, including high osmotic pressure, high temperature, low pH, nutrient deprivation or harmful chemicals (Poger and Mark, 2015).

**Figure 5:**
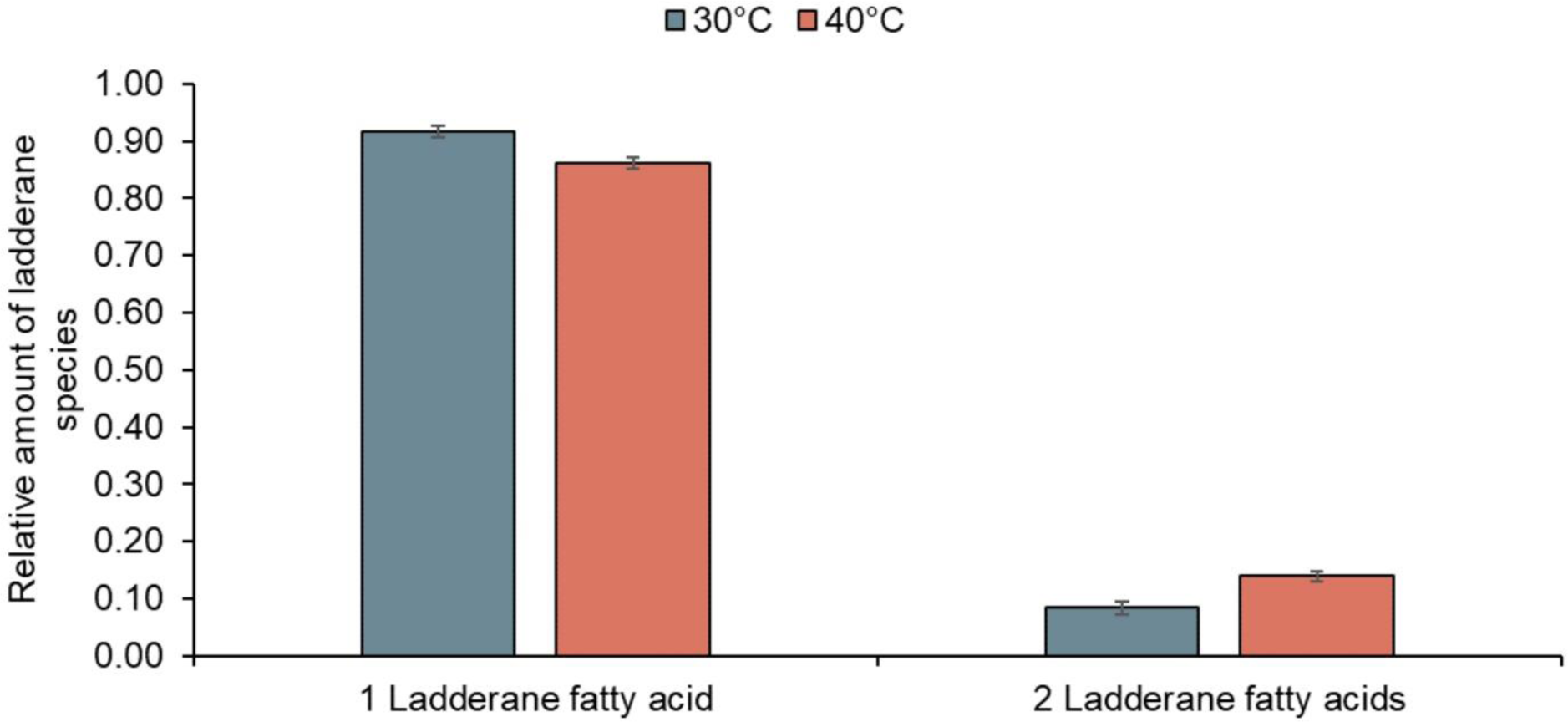
The ladderane cyclization shown as a relative amount of ladderanes with one or both ladderane fatty acids (FAs) at 30 (R0) and 40 °C (R1). The structure of ladderane lipids with one and two ladderane FASs are shown in Fig.4.

Conversely, Koyanagi et al. (2016) and Quehenberger et al. (2020) showed that upon formation of cyclohexane and cyclopentane rings, the leakage of protons slowed and the permeability of membranes reduced when exposed to a higher temperature, lower pH, oxidation, enzymatic degradation, nutrient limitation or generally decreased growth rate.

The major contributor to the almost 1.7-fold higher production of ladderanes with two ladderane FAs was the synthesis of C20 [3]-ladderane FAs, which grew significantly (p-value 0.001), whereas the other ladderane FAs experienced only a mild increase. An increase in this and other ladderane species with temperature is depicted in Fig. 6. C20 [3]-ladderane FA occupied both *sn-1* and *sn-2* positions in a considerable share of membrane consistently across all our four runs without the activity of biomass or reactors performance affecting it. This suggests, that this response is likely similar to the one of other organisms maintaining membrane fluidity. Like other organisms which produce cyclopentane or cyclohexane lipid species, anammox also increased the share of lipids containing cycles with temperature to avoid proton motive force dissipation and not to prevent penetration of hydrazine, supporting the findings of Moss III et al. (2018).

**Figure 6:**
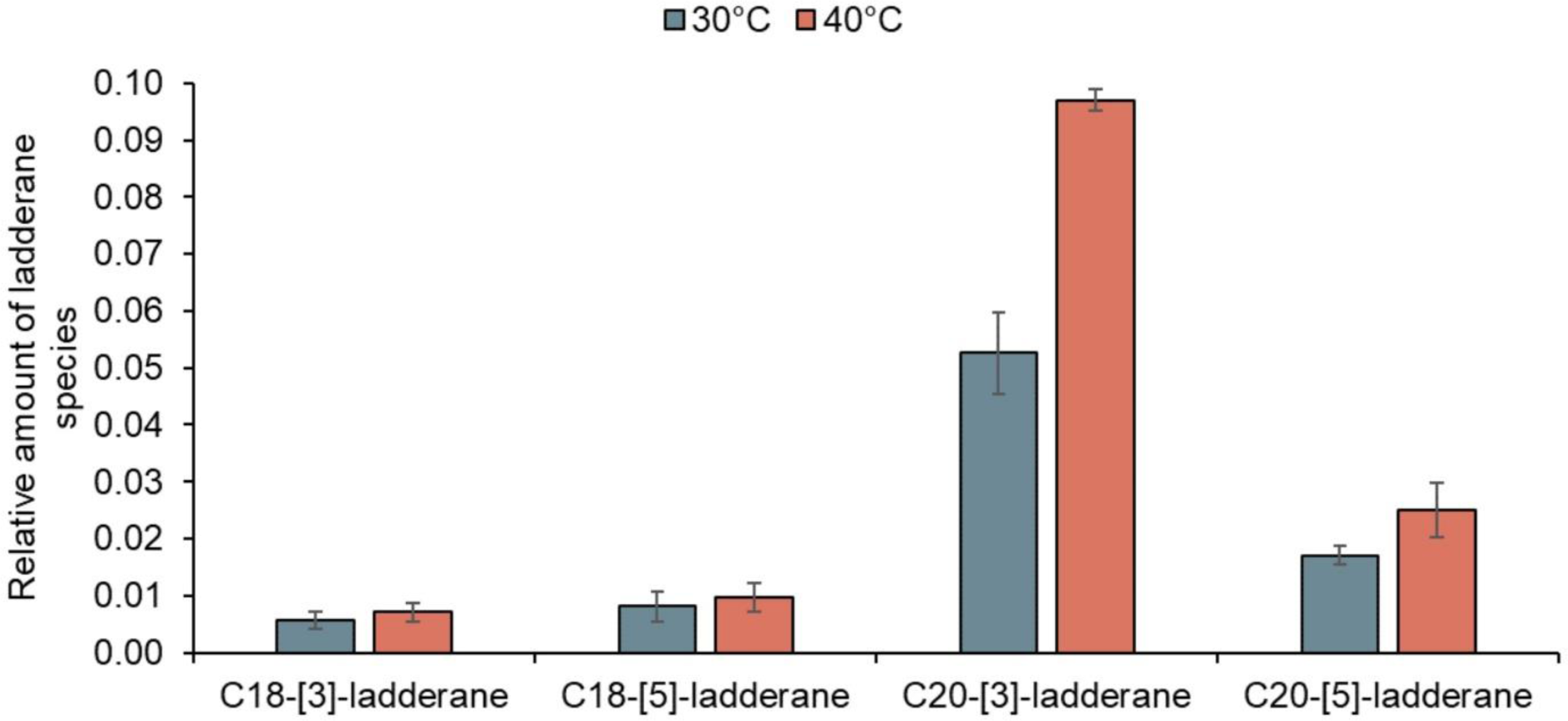
The relative amounts of ladderane fatty acids at 30 (R0) and 40 °C (R1).

Furthermore, this study also showed a slight increase of NL3 (ratio of C20 [3]-ladderanes to the sum of C18 and C20 [3]-ladderanes) and NL5 (identical to NL3 but with [5]-ladderanes) suggested as indicators of anammox adaptation by Kouba et al. (2022c) and Rattray et al. (2010), respectively. The observed growths of NL3 from 0.90±0.01 to 0.93±0.01 and NL5 from 0.69±0.07 to 0.72±0.04 were in line with the previous reports, however, were not found statistically significant in this study. Moreover, the results of Rattray et al. (2010) revealed noticeable changes in NL5 only after more than 20 days and stabilization after 60 days, while R1 in this study was kept at 40 °C for maximally 15 days. Therefore, further verification of this mechanism at thermophilic range over a longer period of time would be beneficial.

#### 3.4.2 Non-ladderane structural alterations

The non-ladderane alkyls composed of either 14, 15 or 16 carbons and comparatively decreased due to the increase in ladderane alkyls. The ratios of non-ladderane C14 alkyls did not change with the temperature as can be seen in Fig. 7. However, the C15 alkyls production rose from 0.37 to 0.50 of the total non-ladderane alkyl chains at thermophilic conditions (p-value 0.008), while C16 decreased from 0.40 to 0.22 (p-value 0.002). These results indicate a shortening of chains rather than their expected elongation with rising temperature, which may present a synergetic mechanism to elevated cyclization. Possibly, C16 alkyls might have been branched and their reduction might further restrict diffusion.

**Figure 7:**
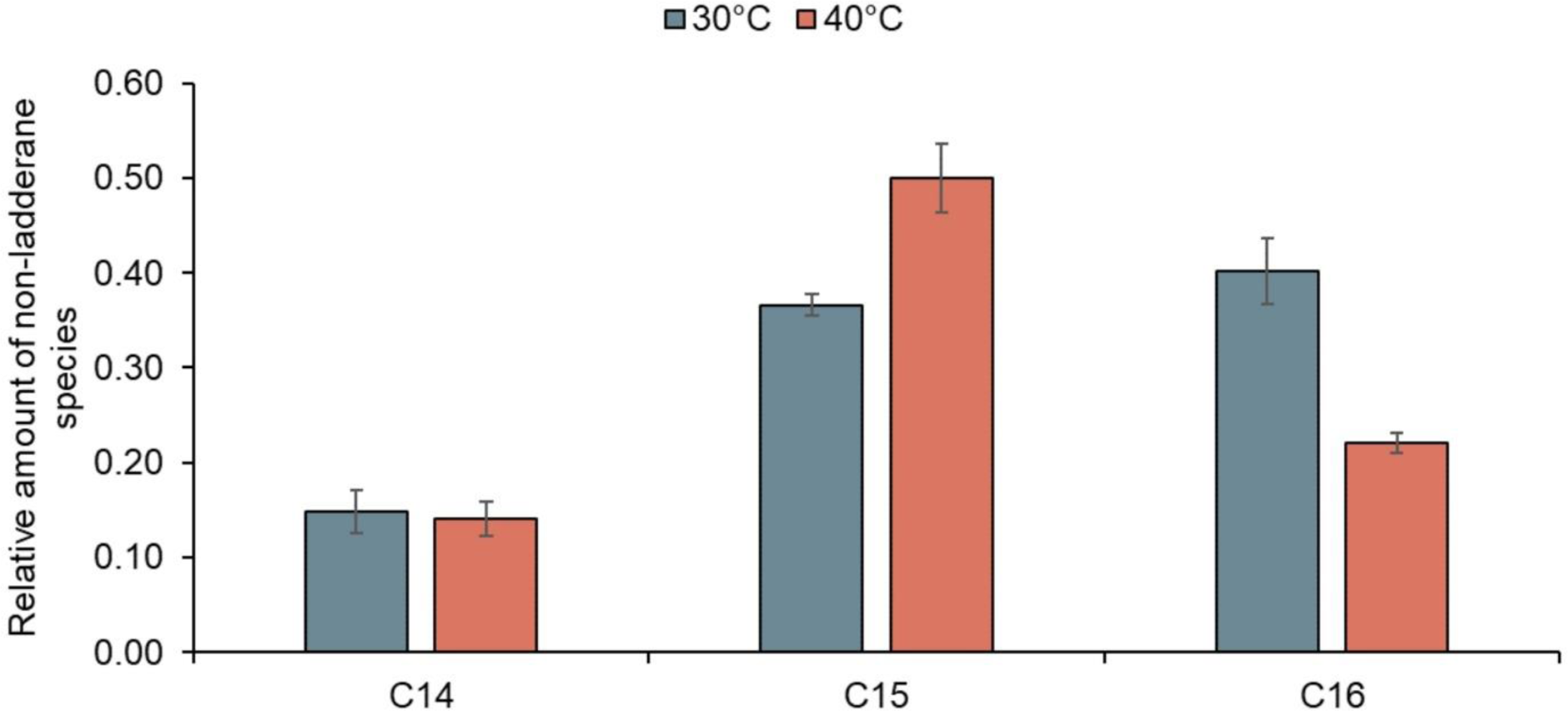
The relative amounts of non-ladderane fatty acids detected in ladderane lipids at 30 (R0) and 40 °C (R1).

In the polar part of lipids, respectively, the relative share of each phosphatidylcholine (FC), phosphatidylethanolamine (FE), or phosphatidylglycerol (FG) did not show a clear trend with increasing temperature varying from 0.21 to 0.55, which is consistent with the results of Kouba et al. (2022c), Kouba et al. (2022b) and Kouba et al. (2022d), as each of these studies found a different polar head group dominating at lower temperatures. Similarly, the ester-to-ether ratio fluctuated from 0.05 to 0.16 without a link to temperature, again correspondingly to the divergencies in previous literature (Kouba et al., 2022c; Rattray et al., 2010). These observed changes suggest an independence on temperature or the effect of factors other than temperature alone.

#### 3.4.3 Discontinuous nature of anammox adaptation via ladderanes

This study found that, under high temperatures, ladderane fatty acids (FAs) in anammox bacteria adapt primarily through changes in cyclization rather than elongation, suggesting two different primary adaptation mechanisms to low and high temperatures.. Siliakus et al. (2017) described discontinuous adaptation mechanisms in other bacteria, namely the cyclopropanes introduction at low temperatures and unsaturation at sub-freezing temperatures. Therefore, we suggest anammox might employ a similar set of various mechanisms proposed in Fig. 8. The adjustment of a number of carbons in the non-ladderane alkyls to adapt to psychrophilic temperatures and cyclization to the thermophilic range. Although the increased cyclization was found in all four runs independent of whether the SAA and NRE were retained or reduced, it still presents an important parameter on anammox viability and adaptation to high temperature conditions. This would be of great impact for water practitioners implementing high-temperature anammox process. Moreover, the discontinuous nature of adaptation is crucial in deducing the anammox distribution, which was previously based on NL5 (van Kemenade et al., 2022).

**Figure 8:**
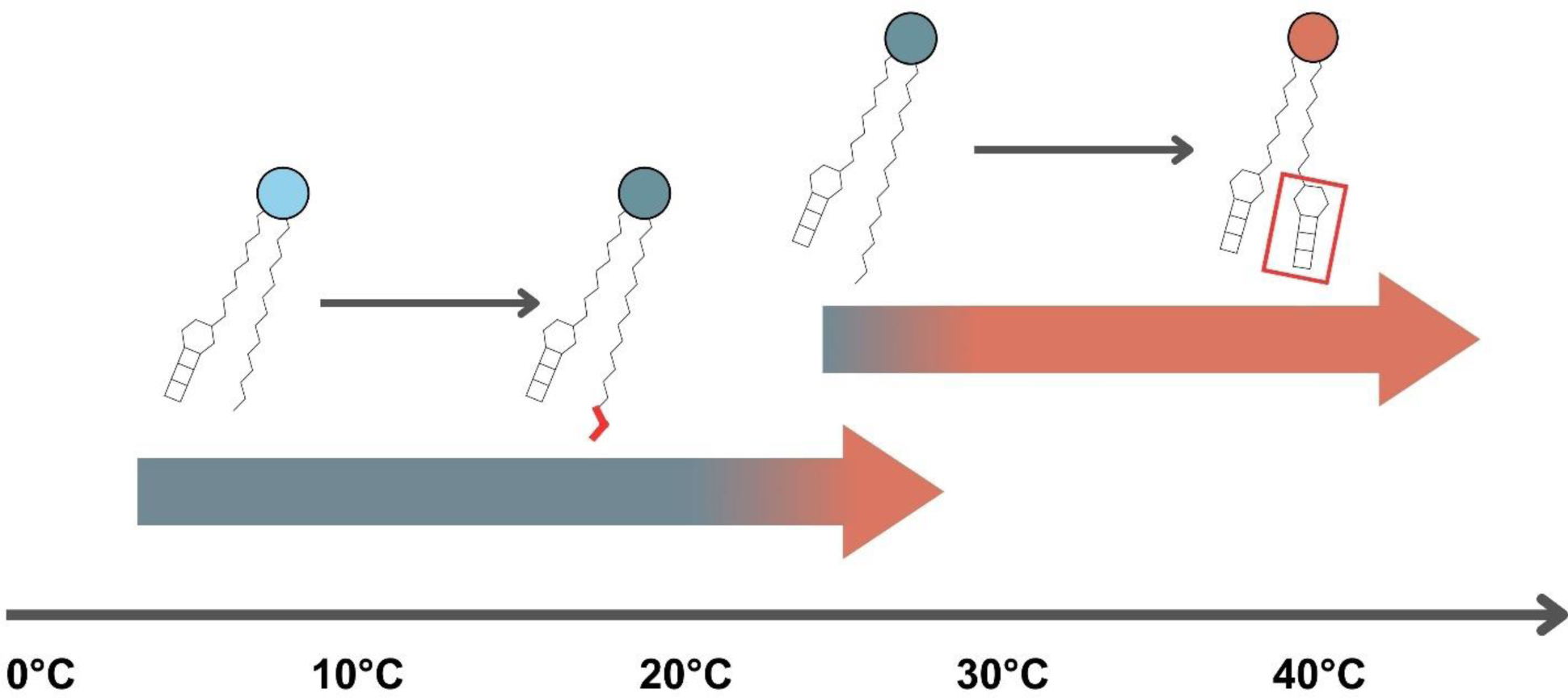
Suggestion of possible discontinuous adaptation via changes in the structure of ladderane lipids (the changes are marked in bright red, first the elongation and then the cyclization). Adaptation up to 30 °C involves adjustment of the length of C14/C15/C16 straight or branched fatty acids and [5]-ladderanes (elongation from C18 to C20, not shown). Adaptation to higher temperatures involves a replacement of straight or branched fatty acids with ladderanes.

Furthermore, we speculate that the observed unprecedented decrease of C16 alkyls in this study not matching the homeoviscous adaptation theory may be the result of ladderanes synthesis, specifically, that C16 alkyls or their parts could be used as an initial substrate for ladderane synthesis by a recently discovered anammox-specific acyl carrier protein and ACP-3-hydroxyacyl dehydratase (Dietl and Barends, 2022; Dietl et al., 2023). However, anammox were found to cope with temperatures as high as 80 °C (Byrne et al., 2009), which implies the existence of another adaptation mechanism to cover these extremely thermophilic temperatures.

### 3.5 Proteome changes under changing temperature

In each of the three runs, LC-MS successfully detected 266, 149 and 157 proteins in both experimental and control biomasses. Of the detected proteins, significant changes were revealed for 1, 18 and 8, respectively (Runs 1-3). Table S2 reveals these changed proteins along with other detected proteins generally involved in the adaptation to high temperatures. Of the proteins generally involved in the temperature adaptation, the anammox bacteria adapted to 40 °C contained slightly higher levels of chaperonin GroEL (protein id: A0A0M2URJ0, A0A0M2UYG9, A0A0M2V1W3). Other proteins generally involved in the high-temperature adaptation as discussed in our previous work Kouba et al. (2022a) were detected but not upregulated, namely GroES which is the co-chaperone to GroEL. We detected also other similar protein chaperones known to co-interact with GroES/GroEL, namely DnaK, ClpB, and HtpG. This applies also to the chaperone PpiC, a homolog of PpiD, a peptide-prolyl isomerase required for folding of outer membrane proteins, thus acting as peri-plasmic chaperone. The increased abundance of chaperones and chaperonins is consistent with the general adaptation mechanisms of microorganisms to increased temperature described by Moon et al. (2023).

Furthermore, we uncovered downregulation patterns consistent with low-temperature adaptation previously described by Lin et al. (2018), Huo et al. (2020) and Wang et al. (2018). These include the downregulation of proteins involved in replication, transcription and translation (DNA gyrase subunit A, Elongation factor G, 30S ribosomal protein, Adenosylhomocysteinase or DNA-directed RNA polymerase subunit beta) and also of the energy production (NADH dehydrogenase, Acetyl-CoA synthase ATPase and ATP synthase). This similarity suggests an adaptation to suboptimal conditions rather than increased/decreased temperature explaining the necessity to halve the SLR when greater temperature adjustments are required.

Moreover, the results of the second run revealed a significant upregulation of 3-oxoacyl-[acyl-carrier-protein] synthase 2, complexed in the FASII system and involved in the lipids biosynthesis (Uegaki et al., 2023). However, its connection with the only described anammox-specific protein speculated to be involved in the ladderane biosynthesis, the anammox-specific acyl carrier protein (ACP) detected in ‘*Ca.* Kuenenia stuttgartiensis’ by Dietl and Barends (2022) is unknown. Importantly, due to the relatively slight changes in the proteome, we suggest that the ladderane biosynthesis responsible for higher ladderane cyclization and RLC was affected on the level of interactome and not on the level of protein synthesis.

To the best of our knowledge, these are the first insights into the adaptation of protein synthesis in anammox bacteria at high temperatures and one of the few studies for bacteria involved in the nitrogen cycle.

## 4 Conclusions

This study is the first to explore the mechanism of anammox adaptation to high temperatures, including the unique ladderane lipids and proteome changes. We showed that high temperature induces downregulation of replication, transcription, translation and energy generation similar to anammox cold-adaptation, negatively influencing the capacity of anammox bacteria to remove nitrogen. However, adjusting specific nitrogen loading below half of the original one can prevent a system deterioration and allows maintaining a successful operation. Surprisingly, anammox bacteria did not adapt to 40 °C via ladderane lipid chain elongation but instead by increasing the number of cyclobutane cycles in ladderane lipids and increasing the content of ladderanes in general not via a protein synthesis but on a level of the interactome. These results suggest, that the introduction of more cyclobutane rings into the ladderane structure presents a novel adaptation mechanism of anammox bacteria towards thermophilic temperatures. Another important mechanism was an upregulation of chaperone GroEL enabling correct protein folding at high temperatures. In conclusion, the adaptation strategies identified in this study present important indicators of anammox adaptation to the conditions of thermophilic anaerobic digestion effluent or industrial wastewaters, emphasizing the imperative for a sufficient adaptation period, particularly as elevated temperatures may pose an even greater challenge than psychrophilic conditions.

## 5 CRediT authorship contribution statement

Karmann, C.: Conceptualization, Formal analysis, Investigation, Methodology, Visualization, Writing – original draft. Navrátilová K.: Formal analysis, Investigation, Methodology. Behner A.: Formal analysis, Investigation. Noor T.: Formal analysis, Investigation. Danner S.: Investigation, Validation. Majchrzak A.: Investigation, Validation. Šantrůček J.: Formal analysis, Investigation, Methodology. Podzimek, T.: Investigation. Marin Lopez, M. A.: Investigation, Data curation. Hajšlová, J.: Methodology, Resources. Bartáček, J.: Funding acquisition, Supervision. Lipovová, P.: Investigation, Data curation, Funding acquisition. Kouba V.: Conceptualization, Project administration, Supervision, Writing - review & editing.

## Acknowledgements

This work was supported by the grant of Specific university research – grant No. A2_FTOP_2022_013 and by the Czech Ministry of Education Youth and Sports through project GACR 17–25781S. The authors also thank to Laura van Niftrik and Guylaine H.L. Nuijten (Microbiology, Radboud University) for providing the inoculum biomass. Moreover, we thank Jana Bartáčková (University of Chemistry and Technology, Prague) for help with analysis of samples and administrative support.

## 7 Data availability

Data will be made available on request.

## References

Apha, A. 1998. Standard methods for the examination of water and wastewater, 20. 632 Washington. DC: american Public health association 633.

Byrne, N., Strous, M., Crépeau, V., Kartal, B., Birrien, J.-L., Schmid, M., Lesongeur, F., Schouten, S., Jaeschke, A. and Jetten, M. 2009. Presence and activity of anaerobic ammonium-oxidizing bacteria at deep-sea hydrothermal vents. The ISME journal 3(1), 117–123.

Chen, Z., Pang, C. and Wen, Q. 2023. Coupled pyrite and sulfur autotrophic denitrification for simultaneous removal of nitrogen and phosphorus from secondary effluent: feasibility, performance and mechanisms. Water Research 243, 120422.

Dietl, A. and Barends, T.R. 2022. Dynamics in an unusual acyl carrier protein from a ladderane lipid-synthesizing organism. Proteins: Structure, Function, and Bioinformatics 90(1), 73–82.

Dietl, A., Wellach, K., Mahadevan, P., Mertes, N., Winter, S.L., Kutsch, T., Walz, C., Schlichting, I., Fabritz, S. and Barends, T.R. 2023. Structures of an unusual 3-hydroxyacyl dehydratase (FabZ) from a ladderane-producing organism with an unexpected substrate preference. Journal of Biological Chemistry 299(5).

Fofana, R., Parsons, M., Long, C., Chandran, K., Jones, K., Klaus, S., Trovato, B., Wilson, C., De Clippeleir, H. and Bott, C. 2022. Full-scale transition from denitrification to partial denitrification– anammox (PdNA) in deep-bed filters: Operational strategies for and benefits of PdNA implementation. Water Environment Research 94(5), e10727.

Fraraccio, S., Strejcek, M., Dolinova, I., Macek, T. and Uhlik, O. 2017. Secondary compound hypothesis revisited: selected plant secondary metabolites promote bacterial degradation of cis-1, 2-dichloroethylene (cDCE). Scientific Reports 7(1), 8406.

Gilbert, E.M., Agrawal, S., Karst, S.M., Horn, H., Nielsen, P.H. and Lackner, S. 2014. Low temperature partial nitritation/anammox in a moving bed biofilm reactor treating low strength wastewater. Environmental science & technology 48(15), 8784–8792.

He, S., Chen, Y., Qin, M., Mao, Z., Yuan, L., Niu, Q. and Tan, X. 2018. Effects of temperature on anammox performance and community structure. Bioresource technology 260, 186–195.

Huang, W., She, Z., Gao, M., Wang, Q., Jin, C., Zhao, Y. and Guo, L. 2019. Effect of anaerobic/aerobic duration on nitrogen removal and microbial community in a simultaneous partial nitrification and denitrification system under low salinity. Science of The Total Environment 651, 859–870.

Huo, T., Zhao, Y., Tang, X., Zhao, H., Ni, S., Gao, Q. and Liu, S. 2020. Metabolic acclimation of anammox consortia to decreased temperature. Environment International 143, 105915.

Iino, T., Mori, K., Uchino, Y., Nakagawa, T., Harayama, S. and Suzuki, K.-i. 2010. Ignavibacterium album gen. nov., sp. nov., a moderately thermophilic anaerobic bacterium isolated from microbial mats at a terrestrial hot spring and proposal of Ignavibacteria classis nov., for a novel lineage at the periphery of green sulfur bacteria. International journal of systematic and evolutionary microbiology 60(6), 1376–1382.

Isaka, K., Date, Y., Kimura, Y., Sumino, T. and Tsuneda, S. 2008. Nitrogen removal performance using anaerobic ammonium oxidation at low temperatures. FEMS microbiology letters 282(1), 32–38.

Jebbar, M., Franzetti, B., Girard, E. and Oger, P. 2015. Microbial diversity and adaptation to high hydrostatic pressure in deep-sea hydrothermal vents prokaryotes. Extremophiles 19(4), 721–740.

Kouba, V., Bachmannová, C., Podzimek, T., Lipovova, P. and Van Loosdrecht, M. 2022a. Physiology of anammox adaptation to low temperatures and promising biomarkers: A review. Bioresource Technology 349, 126847.

Kouba, V., Hůrková, K., Navrátilová, K., Kok, D., Benáková, A., Laureni, M., Vodičková, P., Podzimek, T., Lipovová, P. and van Niftrik, L. 2022b. Effect of temperature on the compositions of ladderane lipids in globally surveyed anammox populations. Science of the Total Environment 830, 154715.

Kouba, V., Hůrková, K., Navrátilová, K., Kok, D., Benáková, A., Laureni, M., Vodičková, P., Podzimek, T., Lipovová, P. and van Niftrik, L. 2022c. On anammox activity at low temperature: Effect of ladderane composition and process conditions. Chemical Engineering Journal 445, 136712.

Kouba, V., Vejmelkova, D., Zwolsman, E., Hurkova, K., Navratilova, K., Laureni, M., Vodickova, P., Podzimek, T., Hajslova, J. and Pabst, M. 2022d. Adaptation of anammox bacteria to low temperature via gradual acclimation and cold shocks: Distinctions in protein expression, membrane composition and activities. Water research 209, 117822.

Koyanagi, T., Leriche, G., Onofrei, D., Holland, G.P., Mayer, M. and Yang, J. 2016. Cyclohexane Rings Reduce Membrane Permeability to Small Ions in Archaea-Inspired Tetraether Lipids. Angewandte Chemie 128(5), 1922–1925.

Li, J., Wang, D., Yu, D. and Zhang, P. 2018. Performance and sludge characteristics of anammox process at moderate and low temperatures. Korean Journal of Chemical Engineering 35, 164–171.

Lin, X., Wang, Y., Ma, X., Yan, Y., Wu, M., Bond, P.L. and Guo, J. 2018. Evidence of differential adaptation to decreased temperature by anammox bacteria. Environmental microbiology 20(10), 3514–3528.

Lotti, T., Kleerebezem, R. and Van Loosdrecht, M. 2015. Effect of temperature change on anammox activity. Biotechnology and bioengineering 112(1), 98–103.

Maiti, A., Kumar, A. and Daschakraborty, S. 2023. How Do Cyclopropane Fatty Acids Protect the Cell Membrane of Escherichia coli in Cold Shock? The Journal of Physical Chemistry B 127(7), 1607–1617.

Moon, S., Ham, S., Jeong, J., Ku, H., Kim, H. and Lee, C. 2023. Temperature matters: bacterial response to temperature change. Journal of Microbiology 61(3), 343–357.

Moss III, F.R., Shuken, S.R., Mercer, J.A., Cohen, C.M., Weiss, T.M., Boxer, S.G. and Burns, N.Z. 2018. Ladderane phospholipids form a densely packed membrane with normal hydrazine and anomalously low proton/hydroxide permeability. Proceedings of the National Academy of Sciences 115(37), 9098–9103.

Narita, Y., Zhang, L., Kimura, Z.-i., Ali, M., Fujii, T. and Okabe, S. 2017. Enrichment and physiological characterization of an anaerobic ammonium-oxidizing bacterium ‘Candidatus Brocadia sapporoensis’. Systematic and applied microbiology 40(7), 448–457.

Nunoura, T., Hirai, M., Miyazaki, M., Kazama, H., Makita, H., Hirayama, H., Furushima, Y., Yamamoto, H., Imachi, H. and Takai, K. 2013. Isolation and characterization of a thermophilic, obligately anaerobic and heterotrophic marine Chloroflexi bacterium from a Chloroflexi-dominated microbial community associated with a Japanese shallow hydrothermal system, and proposal for Thermomarinilinea lacunofontalis gen. nov., sp. nov. Microbes and environments 28(2), 228–235.

Park, G., Takekawa, M., Soda, S., Ike, M. and Furukawa, K. 2017. Temperature dependence of nitrogen removal activity by anammox bacteria enriched at low temperatures. Journal of bioscience and bioengineering 123(4), 505–511.

Pfaffl, M.W. 2001. A new mathematical model for relative quantification in real-time RT–PCR. Nucleic acids research 29(9), e45–e45.

Pfennig, N. and Trüper, H.G. (2019) Handbook of microbiology, pp. 14-24, CRC Press.

Poger, D. and Mark, A.E. 2015. A ring to rule them all: the effect of cyclopropane fatty acids on the fluidity of lipid bilayers. The journal of physical chemistry B 119(17), 5487–5495.

Quehenberger, J., Pittenauer, E., Allmaier, G. and Spadiut, O. 2020. The influence of the specific growth rate on the lipid composition of Sulfolobus acidocaldarius. Extremophiles 24, 413–420.

Rattray, J.E., van de Vossenberg, J., Jaeschke, A., Hopmans, E.C., Wakeham, S.G., Lavik, G., Kuypers, M.M., Strous, M., Jetten, M.S. and Schouten, S. 2010. Impact of temperature on ladderane lipid distribution in anammox bacteria. Applied and Environmental Microbiology 76(5), 1596–1603.

Ribeiro, H., Wijaya, I.M.W., Soares-Santos, V., Soedjono, E.S., Slamet, A., Teixeira, C. and Bordalo, A.A. 2022. Microbial community composition, dynamics, and biogeochemistry during the start-up of a partial nitritation-anammox pathway in an upflow reactor. Sustainable Environment Research 32(1), 18.

Siliakus, M.F., van der Oost, J. and Kengen, S.W. 2017. Adaptations of archaeal and bacterial membranes to variations in temperature, pH and pressure. Extremophiles 21, 651–670.

Sobotka, D., Czerwionka, K. and Makinia, J. 2016. Influence of temperature on the activity of anammox granular biomass. Water Science and Technology 73(10), 2518–2525.

Sobotka, D., Zhai, J. and Makinia, J. 2021. Generalized temperature dependence model for anammox process kinetics. Science of The Total Environment 775, 145760.

Uegaki, T., Takei, T., Yamaguchi, S., Fujiyama, K., Sato, Y., Hino, T. and Nagano, S. 2023. Anammox Bacterial S-Adenosyl-l-Methionine Dependent Methyltransferase Crystal Structure and Its Interaction with Acyl Carrier Proteins. International Journal of Molecular Sciences 24(1), 744.

van Kemenade, Z.R., Villanueva, L., Hopmans, E.C., Kraal, P., Witte, H.J., Sinninghe Damsté, J.S. and Rush, D. 2022. Bacteriohopanetetrol-x: constraining its application as a lipid biomarker for marine anammox using the water column oxygen gradient of the Benguela upwelling system. Biogeosciences 19(1), 201–221.

Vandekerckhove, T.G., Props, R., Carvajal-Arroyo, J.M., Boon, N. and Vlaeminck, S.E. 2020. Adaptation and characterization of thermophilic anammox in bioreactors. Water Research 172, 115462.

Wang, W., Yan, Y., Song, C., Pan, M. and Wang, Y. 2018. The microbial community structure change of an anaerobic ammonia oxidation reactor in response to decreasing temperatures. Environmental Science and Pollution Research 25, 35330–35341.

Wang, X.-W., Tan, X., Dang, C.-C., Lu, Y., Xie, G.-J. and Liu, B.-F. 2023. Thermophilic microorganisms involved in the nitrogen cycle in thermal environments: Advances and prospects. Science of the Total Environment, 165259.

Yuan, Q., Wang, K., He, B., Liu, R., Qian, L., Wan, S., Zhou, Y., Cai, H. and Gong, H. 2021. Spontaneous mainstream anammox in a full-scale wastewater treatment plant with hybrid sludge retention time in a temperate zone of China. Water Environment Research 93(6), 854–864.

Zhang, Z.-Z., Ji, Y.-X., Cheng, Y.-F. and Jin, R.-C. 2018. Increased salinity improves the thermotolerance of mesophilic anammox consortia. Science of the total environment 644, 710–716.

Zhu, W., Li, J., Dong, H., Wang, D. and Zhang, P. 2017. Nitrogen removal performance and operation strategy of anammox process under temperature shock. Biodegradation 28, 261–274.

