## Supplementary materials for "Mechanisms of Anammox Adaptation to High Temperatures: Increased Cyclization of Ladderane Lipids and Proteomic Insights"

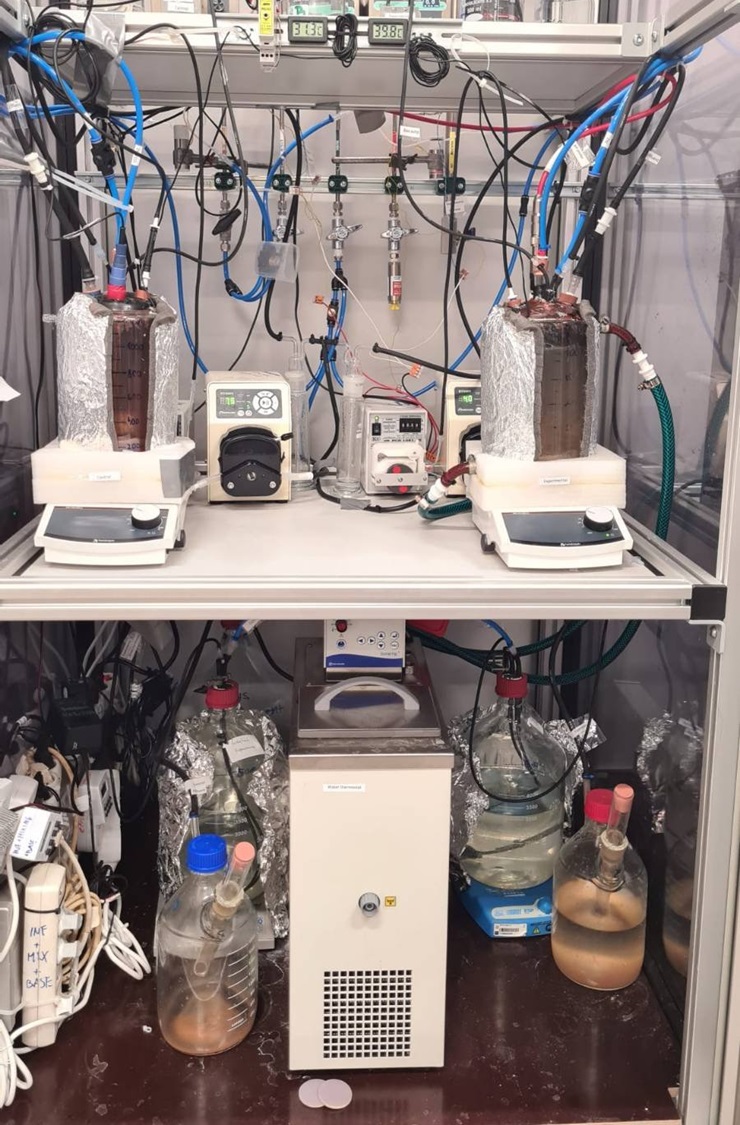

**Figure S *1*:** Photography of the experimental setup.

**Table S *1*:** Composition of the mineral medium / feed to the reactors.

| **Compound** | **Concentration (mg L^-1^)** |
| --- | --- |
| Calcium chloride (CaCl_2_) | 100 |
| Magnesium Sulphate (MgSO_4_) | 300 |
| Potassium Hydrogen Phosphate (KH_2_PO_4_) | 30 |
| Potassium Hydrogen Carbonate (KHCO_3_) | 500 |
| Iron Sulphate (FeSO_4_.7H_2_O) | 5 |
| Hydrogen Chloride (HCl) | 1 |
| Trace Elements | 1^a, b^ |

a. Unit: mL L^-1^
b. Trace elements composition was taken from Van de Graaf et al. (1995)

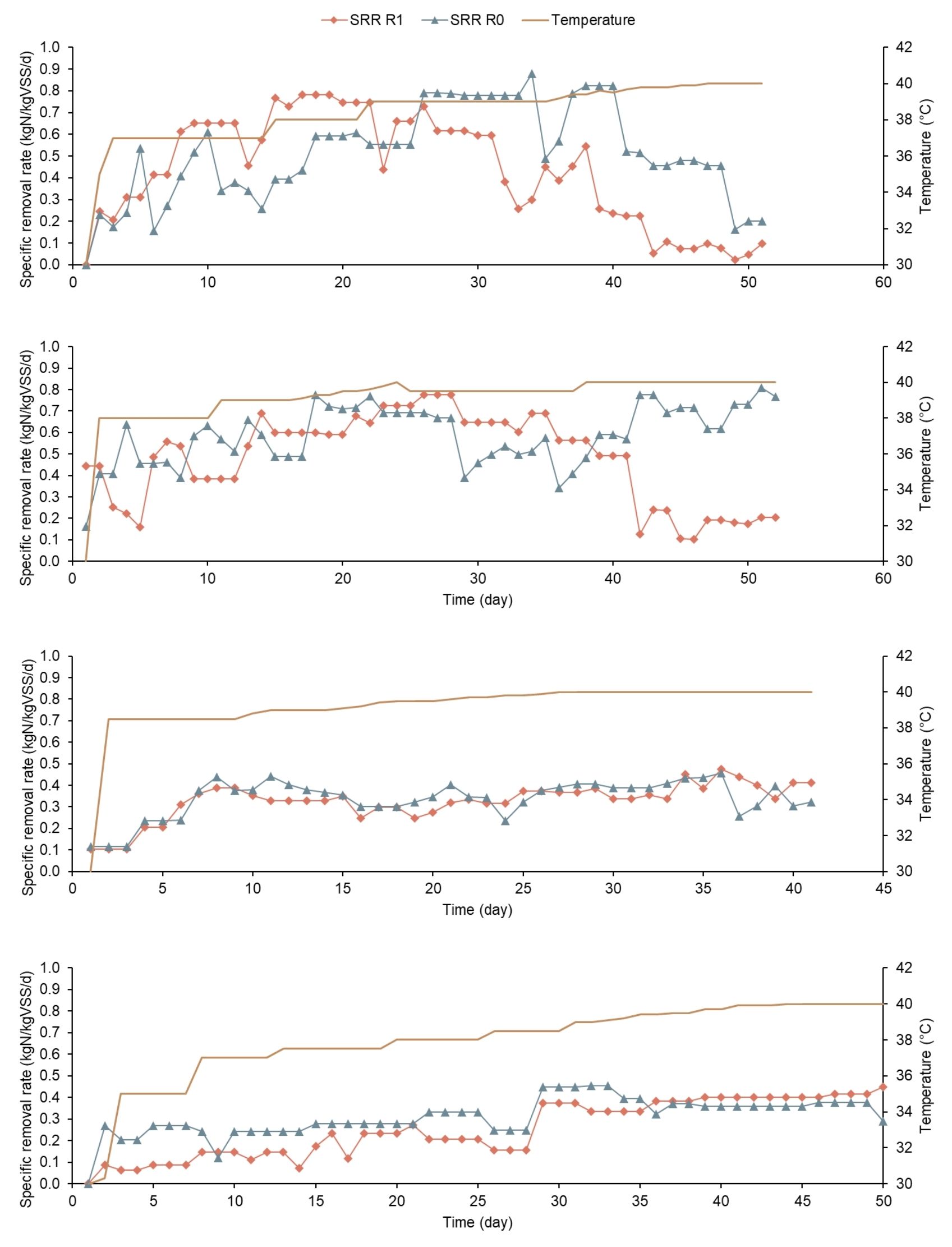

**Figure S *2*:** Development of specific nitrogen removal rate (SRR) of R0 and R1 and temperature in the four Runs. A) Run 1, B) Run 2, C) Run 3, D) Run 4.

**Table S *2*:** The proteins detected in Runs 2-4. The table shows the results of proteomics, including the protein description, gene tag, ID, relative content and change and also whether the change was statistically significant (p < 0.05, F>F_crit_).

| **Run 2** |  | | | |  |  |  |
| --- | --- | --- | --- | --- | --- | --- | --- |
| **Description, gene tag** | **Relative content (log2)**  **40 °C** | | | **Relative content (log2)**  **30 °C** | **Relative change (%)** | **ANOVA-significant** | **Protein ID** |
| Chaperonin GroEL, groEL | 23.2 | | | 22.7 | 145 | - | A0A0M2URJ0 |
| Chaperone protein DnaK, dnaK | 23.9 | | | 23.6 | 124 | - | A0A0M2US87 |
| Universal stress protein, BROFUL_02478 | 18.7 | | | 20.6 | 26 | - | A0A0M2USK1 |
| Chaperone protein ClpB, clpB | 21.2 | | | 20.9 | 123 | - | A0A0M2UTM1 |
| Universal stress protein, BROFUL_02090 | 19.4 | | | 19.5 | 94 | - | A0A0M2UTS2 |
| DNA gyrase subunit A, gyrA | 18.4 | | | 19.0 | 66 | - | A0A0M2UU87 |
| PpiC domain-containing protein, BROFUL_02831 | 21.4 | | | 20.1 | 236 | - | A0A0M2UVE7 |
| Chaperonin GroEL, groEL | 23.1 | | | 22.7 | 131 | - | A0A0M2UY55 |
| Chaperonin GroEL, groEL | 23.9 | | | 23.6 | 122 | - | A0A0M2UYG9 |
| Chaperonin GroEL, groEL | 23.5 | | | 23.1 | 135 | - | A0A0M2V1W3 |
| Chaperone protein HtpG, htpG | 21.9 | | | 21.9 | 99 | - | A0A0M2V2R4 |
| **Run 3** |  | | |  |  |  |  |
| **Description, gene tag** | **Relative content (log2)**  **40 °C** | **Relative content (log2)**  **30 °C** | | | **Relative change (%)** | **ANOVA-significant** | **Protein ID** |
| Uncharacterized protein, GN=BROFUL_02819 | 22.2 | | | 20.5 | 330 | + | A0A0M2US75 |
| Elongation factor G, fusA | 19.4 | | | 20.8 | 40 | + | A0A0M2USI9 |
| Thioredoxin, BROFUL_01877 | 20.3 | | | 22.1 | 29 | + | A0A0M2UUW5 |
| Polyribonucleotide nucleotidyltransferase, pnp | 19.0 | | | 20.6 | 34 | + | A0A0M2UV21 |
| Methionine adenosyltransferase, BROFUL_01814 | 21.5 | | | 22.4 | 56 | + | A0A0M2UV60 |
| dTDP-glucose 4,6-dehydratase, BROFUL_02837 | 18.1 | | | 19.2 | 49 | + | A0A0M2UVC3 |
| NADH oxidase, BROFUL_02747 | 18.5 | | | 19.6 | 48 | + | A0A0M2UVM2 |
| Glutamine--fructose-6-phosphate aminotransferase [isomerizing], glmS | 18.2 | | | 19.2 | 50 | + | A0A0M2UWG7 |
| 30S ribosomal protein S1, BROFUL_01035 | 19.4 | | | 21.3 | 27 | + | A0A0M2UWI0 |
| Adenosylhomocysteinase, ahcY | 21.9 | | | 23.4 | 36 | + | A0A0M2UWX9 |
| ABC transporter ATP-binding component, BROFUL_01623 | 22.1 | | | 21.0 | 211 | + | A0A0M2UXK1 |
| 3-oxoacyl-[acyl-carrier-protein] synthase 2, BROFUL_01969 | 20.9 | | | 20.0 | 184 | + | A0A0M2UXW4 |
| NADH dehydrogenase, BROFUL_00754 | 21.5 | | | 22.7 | 43 | + | A0A0M2UY70 |
| 30S ribosomal protein S2, rpsB | 20.9 | | | 21.9 | 48 | + | A0A0M2UYU9 |
| Putative flavoprotein, BROFUL_00143 | 19.5 | | | 18.6 | 178 | + | A0A0M2UZA6 |
| Transcription termination factor Rho, rho | 20.0 | | | 21.1 | 46 | + | A0A0M2UZU3 |
| Aspartate--tRNA(Asp/Asn) ligase, aspS | 21.4 | | | 20.9 | 143 | + | A0A0M2V269 |
| Putative heat shock protein, BROFUL_00037 | 18.7 | | | 20.4 |  | + | A0A0M2UZI2 |
| Chaperonin GroEL, groEL | 23.1 | | | 22.6 | 31 | - | A0A0M2URJ0 |
| Chaperone protein DnaK, dnaK | 23.2 | | | 23.1 |  | - | A0A0M2US87 |
| Universal stress protein, BROFUL_02478 | 20.0 | | | 19.7 | 138 | - | A0A0M2USK1 |
| Chaperone protein ClpB, clpB | 20.9 | | | 20.9 | 107 | - | A0A0M2UTM1 |
| PpiC domain-containing protein, BROFUL_02831 | 20.4 | | | 19.6 | 123 | - | A0A0M2UVE7 |
| Cold shock protein, BROFUL_01715 | 19.8 | | | 21.1 | 96 | - | A0A0M2UX69 |
| Chaperonin GroEL, groEL | 22.9 | | | 22.7 | 169 | - | A0A0M2UY55 |
| Chaperonin GroEL, groEL | 23.6 | | | 23.4 | 42 | - | A0A0M2UYG9 |
| Chaperonin GroEL, groEL | 23.8 | | | 23.6 | 108 | - | A0A0M2V1W3 |
| **Run 4** |  | | |  |  |  |  |
| **Description, gene tag** | **Relative content**  **(log2)**  **40 °C** | | **Relative content**  **(log2)**  **30 °C** | | **Relative change (%)** | **ANOVA-significant** | **Protein ID** |
| Acetyl-CoA synthase, BROFUL_02730 | 18.5 | | | 19.2 | 62 | + | A0A0M2URU3 |
| DNA-directed RNA polymerase subunit beta, rpoC | 21.1 | | | 21.9 | 59 | + | A0A0M2UTT9 |
| Peptidase, BROFUL_02231 | 20.3 | | | 21.5 | 43 | + | A0A0M2UTW6 |
| ATPase , BROFUL_02791 | 19.8 | | | 20.7 | 51 | + | A0A0M2UU51 |
| ATP synthase subunit beta, atpD | 19.7 | | | 20.2 | 71 | + | A0A0M2UVI2 |
| Chaperonin GroEL, groEL | 24.8 | | | 23.1 |  | + | A0A0M2URJ0 |
| Chaperonin GroEL, groEL | 25.6 | | | 24.1 | 329 | + | A0A0M2UYG9 |
| Chaperonin GroEL, groEL | 25.8 | | | 24.3 | 289 | + | A0A0M2V1W3 |
| Chaperone protein DnaK, dnaK | 24.9 | | | 24.6 | 278 | - | A0A0M2US87 |
| Chaperone protein ClpB, clpB | 22.2 | | | 22.1 |  | - | A0A0M2UTM1 |
| DNA gyrase subunit A, GyrA | 18.6 | | | 19.1 | 126 | - | A0A0M2UU87 |
| PpiC domain-containing protein | 22.0 | | | 21.5 | 109 | - | A0A0M2UVE7 |
| Chaperonin GroEL, groEL | 24.7 | | | 23.6 | 69 | - | A0A0M2UY55 |
| Co-chaperonin GroES, groES | 23.1 | | | 20.6 | 146 | - | A0A0M2UZ82 |
| Chaperone protein HtpG, htpG | 22.6 | | | 22.8 | 212 | - | A0A0M2V2R4 |

1. Present affiliation: Energieinstitut at the Johannes Kepler University in Linz, Department of Energy Technology, Altenberger Straße 69, 4040 Linz, Austria
2. Present affiliation:
